# Inhibitory gating of thalamocortical inputs onto rat gustatory insular cortex

**DOI:** 10.1101/2022.12.08.519650

**Authors:** Melissa S. Haley, Alfredo Fontanini, Arianna Maffei

**Author notes:** **Data Availability.** All data needed to evaluate the conclusions in the paper are present in the paper and/or the Supplementary Materials. **Author Contributions:** Designed experiments: MSH, AF, AM Performed experiments: MSH Analyzed data: MSH Supervised experiments and analysis: AF, AM Writing: MSH, AF, AM.

## Abstract

In the rat primary gustatory cortex (GC), a subregion of the larger insular cortex, neurons display time-varying neural responses to gustatory stimuli. GC taste responses are dramatically reduced following inactivation of the gustatory thalamus, the parvicellular region of the ventral posteromedial thalamic nucleus (VPMpc). Pharmacological inactivation of VPMpc also has a profound effect on GC spontaneous activity. This indicates that the projection from VPMpc plays a crucial role in GC taste processing as well as in the control of its state. How VPMpc afferents engage GC circuits and drive neuronal ensembles to effectively code tastant identity, as well as modulate the overall state of the GC network, remains unclear. To investigate the synaptic properties and organization of VPMpc afferents in GC, we employed a circuit-breaking optogenetic approach, stimulating VPMpc terminal fields while performing whole-cell patch clamp recordings from GC neurons in rat acute slices. Informed by previous studies of thalamocortical inputs to other sensory cortices, we hypothesized that VPMpc-GC synapses have laminar- and cell-specific properties that gate sensory input, conferring computationally flexibility to how taste information is processed in GC. We found that VPMpc-GC synapses are strongly gated by the activity regime of VPMpc afferents, as well as by feedforward and feedback inhibition onto VPMpc terminals. These results provide novel insight into the circuit underpinning of GC responsiveness to incoming thalamocortical activity.

**SIGNIFICANCE STATEMENT:** We report that the input from the primary taste thalamus to the primary gustatory cortex (GC) shows distinct properties compared to primary thalamocortical synapses onto other sensory areas. VPMpc afferents in GC make synapses with excitatory neurons distributed across all cortical layers and display frequency-dependent short-term plasticity to repetitive stimulation, thus they do not fit the classic distinction between drivers and modulators typical of other sensory thalamocortical circuits. Feedforward inhibition gates thalamocortical activation and provides local corticothalamic feedback via presynaptic ionotropic and metabotropic GABA receptors. The connectivity and inhibitory control of thalamocortical synapses support the time-varying response dynamics to taste stimuli observed in GC neurons *in vivo*.

## INTRODUCTION

The rat gustatory cortex (GC), located within the larger insular cortex, is the primary cortical area for the processing of taste and taste-related information. In response to tastant delivery, GC neurons produce time-varying changes in spiking activity that encode both chemosensory and affective variables associated with taste (Katz et al., 2001; Jones et al., 2007). Studies *in vivo* revealed that these spiking dynamics arise via input from two key pathways to GC: the thalamocortical, sensory pathway via the parvicellular portion of the ventral posteromedial nucleus of the thalamus (gustatory thalamus, VPMpc)(Pritchard et al., 1989; Verhagen et al., 2003; Samuelsen et al., 2013), and the limbic pathway, via the basolateral nucleus of the amygdala (BLA) (Piette et al., 2012; Lin et al., 2021). Pharmacological inactivation of VPMpc or BLA in naïve rats greatly affects GC neural activity related to taste quality and palatability (Piette et al., 2012; Samuelsen et al., 2012; Samuelsen et al., 2013). In addition, silencing of the VPMpc dramatically affect spontaneous activity in GC, demonstrating a role in maintaining and modulating the overall state of gustatory cortical networks (Samuelsen et al., 2013).

While we have some understanding of the synaptic organization of the BLA-GC input (Haley et al., 2016), very little is known regarding VPMpc-GC connectivity and how thalamocortical input recruits GC circuits. In other primary sensory cortices, thalamocortical inputs exhibit both laminar- and cell-type specific properties that can strongly inform function and impose patterns of activation dependent on specific connectivity motifs. For example, in visual cortex, thalamocortical afferents onto Layer 4 neurons strongly drive the circuit, whereas projections to Layer 6 are sparser and weaker, and modulate circuit activity (Olsen et al., 2012; Wang et al., 2013). Furthermore, there is evidence that thalamocortical projections directly target inhibitory neurons, driving feedforward inhibition (Gabernet et al., 2005; Cruikshank et al., 2007). This feedforward inhibition can serve multiple roles such as delimiting temporal windows for the integration of diverse inputs (Wehr and Zador, 2003), modulating the gain of excitatory neurons (Cardin et al., 2008), modifying the signal to noise ratio and generating oscillatory activity. In GC, *in vivo* intracellular recordings in anesthetized animals demonstrated that electrical stimulation of VPMpc resulted in two types of synaptic responses in GC neurons, distinguished by the presence or absence of an inhibitory component. This VPMpc-evoked polysynaptic inhibition greatly affected how neurons temporally integrated VPMpc and limbic inputs (Stone et al., 2020).

In the present work, we sought to identify the connectivity, postsynaptic targets and synaptic properties of thalamocortical projections from VPMpc to GC and investigated how VPMpc afferents engage local GABAergic circuits to influence synaptic dynamics. Differently from other sensory cortices, we found that VPMpc afferents to GC target excitatory neurons across all layers. However, laminar specificity could be detected by comparing amplitudes of evoked responses. In addition, we unveiled unique frequency-dependent short-term dynamics of VPMpc-GC evoked responses. Finally, we demonstrate that VPMpc afferents recruit local inhibitory circuits in GC and engage both feedforward inhibition onto postsynaptic neurons and feedback inhibition onto VPMpc axon terminals, pointing at GC inhibitory drive as a gating mechanism modulating incoming activity. Together these data provide insight into the mechanisms by which a sensory thalamic input can engage cortical circuits to enable both temporal coding of taste information as well as global changes in network state.

## METHODS

Long Evans rats of both sexes were used for this study. All animals were group housed on a 12h/12h light dark cycle and received ad libitum access to food and water. All experiments and surgical procedures were approved by the Stony Brook University IACUC and followed the National Institute of Health guidelines.

### Viral injections

Male and female rats aged P14 were anesthetized with a ketamine/xylazine/acepromazine cocktail via intraperitoneal injection (70mg/kg ketamine, 0.7mg/kg acepromazine, and 3.5mg/kg xylazine). Animals were mounted in a stereotaxic frame and a craniotomy was opened at -2.8mm from bregma and 1.0mm from midline to target injection to VPMpc (6.3 dorsal/ventral). A Drummond nanoject was used to inject 50nl of AAV9.CAG.ChR2-Venus.WPRE.SV40 construct (Petreanu et al., 2007) containing 5.64^12^ particles/mL at a rate of 0.5nl/s. Animals recovered for 14 days to allow for full expression of the virus throughout the entirety of VPMpc axonal projections to GC before slice recordings were performed. Slices containing VPMpc were obtained to visually confirm accuracy of injection placement by imaging the GFP co-expression and consistency of viral expression across animals was assessed using a calibration curve as previously described (Haley et al., 2016).

### Slice electrophysiology

Following surgery recovery and viral incubation period, acute slices containing GC were obtained. Animals were anesthetized with isofluorane (bell-jar to effect) and quickly decapitated. Brains were dissected in ice-cold ACSF and then 300μm coronal slices containing GC (+1.5mm to bregma) were obtained on a vibratome (Leica VT1000). Slices were heated to 35ºC in a hot water bath and then maintained at room temperature for the remainder of the experiment. ACSF contained the following (mM): 126 NaCl, 3 KCl, 25 NaHCO3, 1 NaHPO4, 2 MgSO4, 2 CaCl2, 14 dextrose with an osmolarity of 315-325mOsm. For all experiments except Fig. 4, the internal solution contained (mM): 116 K-Glu, 4 KCl, 10 K-HEPES, 4 Mg-ATP, 0.3 Na-GTP, 10 Na-phosphocreatine, 0.4% biocytin (V_rev_ [Cl^−1^] = −81mV). The internal solution to isolate the excitatory and inhibitory components of the VPMpc evoked response (Fig. 4) contained (mM): 20 KCl, 100 Cs-sulfate, 10 K-HEPES, 4 Mg-ATP, 0.3 Na-GTP, 10 Na-phosphocreatine, 3 QX-314 (Tocris Bioscience), 0.2% biocytin (Vrev [Cl^−1^] = −49.3mV). Internal solutions were adjusted to a pH of 7.35 with KOH and an osmolarity of 295mOsm with sucrose. To assess the contribution of cortical GABA to the VPMpc-evoked response, the following GABA agonists where bath applied sequentially and additively: 1mM muscimol (Tocris Bioscience), 10μM baclofen (Tocris Bioscience). Virally transfected VPMpc afferents were activated with 5ms 470nm light pulses (4.4mW) controlled with an LED driver (ThorLabs) and delivered through the 40x objective mounted on an upright microscope (Olympus BX51WI).

### Data analysis

Data were analyzed using Excel, GraphPad Prism, and custom procedures in Igor. For two-group comparisons we used estimation statistics (Ho et al., 2019) to calculate the 95% confidence interval of the mean difference via bootstrap resampling (5000 samples) and included histograms of the resampling distribution of the difference in the means in the figures. For multiple group comparisons we compared the mean of each group to L4 (Fig. 1) or the control (Fig. 5D). For two-group comparisons with multiple conditions (i.e., stimulation frequency, drugs) we used 2-way ANOVAs with post-hoc paired or unpaired permutation t-tests or post-hoc Sidak’s or Tukey’s multiple comparisons tests, as appropriate. Chi-square tests were used to assess VPMpc connectivity, and the proportion of facilitating/depressing synapses. Spearman rank order correlation was used to compare Δ holding current to Δ VPMpc-evoked excitatory postsynaptic currents (VPMpc-EPSC) amplitude in the drug experiments. Cortical depth ratio was calculated as the distance of the recorded neuron from pia divided by the distance between pia and the white matter. All data are presented as the mean ± SEM.

**Figure 1:**
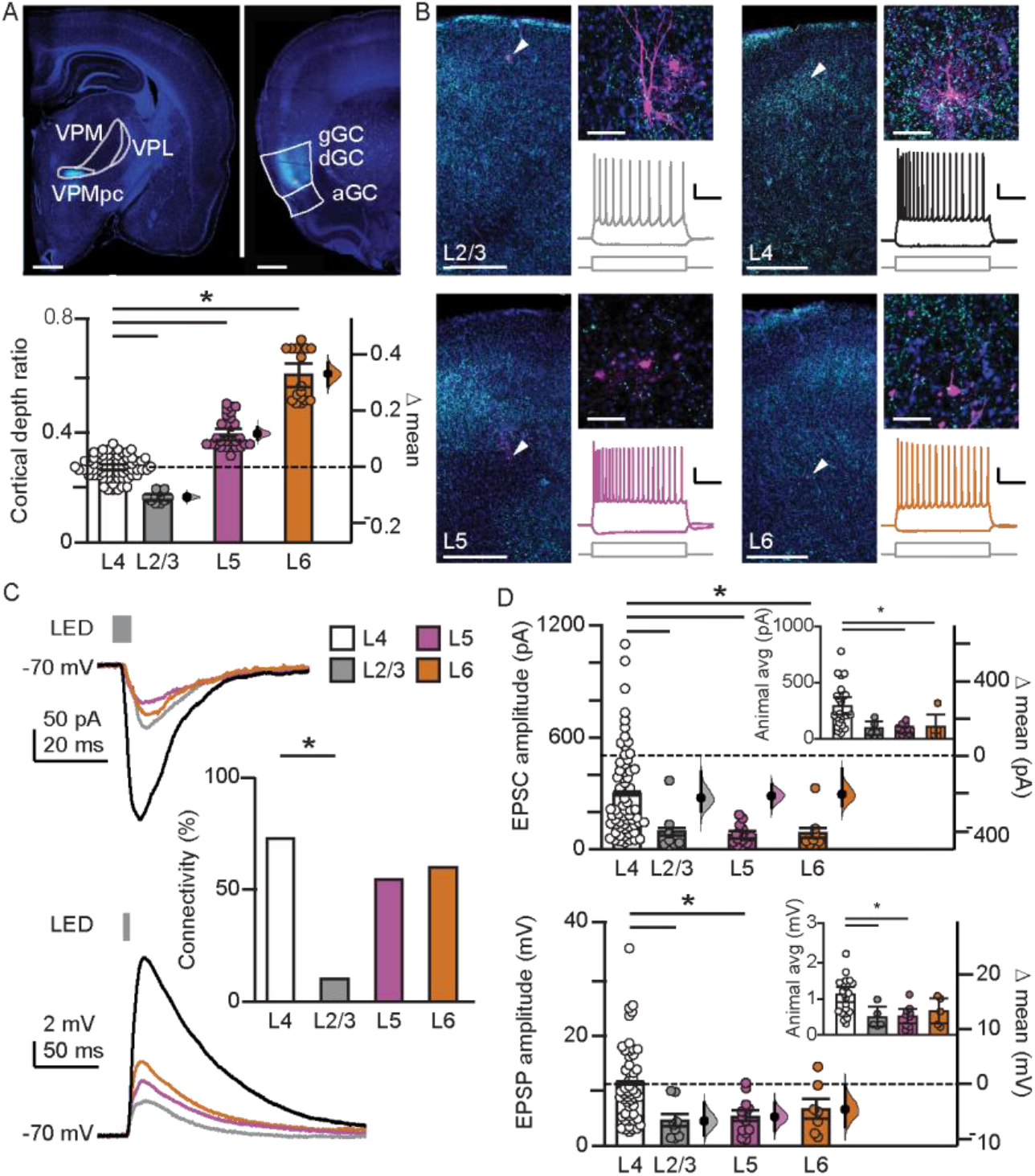
VPMpc afferent input to GC has laminar-specific properties. **A**. Top: Representative images of injection site in rat VPMpc (left) and VPMpc terminal fields in GC (right); blue: Hoescht, cyan: CAG-AAV9-eYFP-ChR2. Scale bar = 1mm. Bottom: Summary of cortical depth ratio (distance of neuron from pia/distance of pia from white matter) for neurons recorded in L2/3 (n=11), L4 (n=73), L5 (n=28), and L6 (n=17). **B**. Representative images of laminar location of biocytin-filled neurons and sample traces of firing pattern to steady-state current injection (−50pA, 250pA) for pyramidal neurons recorded in L2/3 (top left), L4 (top right), L5 (bottom left), and L6 (bottom right). Blue: Hoescht; cyan: CAG-AAV9-eYFP-ChR2; magenta: biocytin. Scale bar, laminar location = 250um, biocytin fill = 50um, firing pattern = 20mV/200ms. **C**. Top: Sample traces of VPMpc-EPSCs onto L2/3, L4, L5, and L6 excitatory neurons recorded at -70 mV. Middle: Sample traces of VPMpc-EPSPs onto L2/3, L4, L5, and L6 excitatory neurons recorded at -70 mV. Bottom: Summary of connectivity percentage across layers (L2/3, n=10/21; L4, n=87/119; L5, n=17/31; L6, n=12/19). **D**. Top: Summary of VPMpc-EPSC amplitude on L2/3 (n=7), L4 (n=67), L5 (n=14), and L6 (n=8) excitatory neurons and mean difference from L4 (Inset: Average EPSC amplitude by animal: L2/3, n=5; L4, n=24; L5, n=10; L6, n=6). Bottom: Summary of VPMpc-EPSP amplitude on L2/3 (n=7), L4 (n=53), L5 (n=7), and L6 (n=6) L6 excitatory neurons and mean difference from L4 (Inset: Average EPSP amplitude by animal: L2/3, n=5; L4, n=23; L5, n=10; L6, n=5). * indicates p ≥ 0.05. Error bars ± SEM.

### Immunohistochemistry

After recording, acute slices were post-fixed in 4% PFA for 1-2 weeks. They were then washed in PBS (3 × 10min rinse), permeabilized and blocked in a solution containing 1% Triton X and 10% normal goat serum in PBS for 2h, then incubated overnight at 4°C in a solution containing 0.1% Triton X and 3% normal goat serum in PBS, streptavidin Alexa Fluor-568 conjugate (1:2000, Invitrogen, S11226), mouse anti-GAD67 (1:500, MilliporeSigma, MAB5406, monoclonal), and chicken anti-GFP (1:1000, Abcam, ab13970, polyclonal). GC slices were then rinsed in PBS (3 × 10min) and incubated at 25°C for 2h in a solution containing 0.1% Triton X and 3% normal goat serum in PBS, goat anti-mouse Alexa Fluor-647 (1:500, Invitrogen, A21235), goat anti-chicken Alexa Fluor-488 (1:500, Abcam, ab150173), and Hoechst 33342 (1:5000, Invitrogen, H3570). Slices containing VPMpc were incubated overnight at 4°C in a solution containing 0.1% Triton X and 3% normal goat serum in PBS and chicken anti-GFP (1:1000, Abcam, ab13970, polyclonal). They were then rinsed in PBS (3 × 10min) and incubated at 25°C for 2h in a solution containing 0.1% Triton X and 3% normal goat serum in PBS, goat anti-chicken Alexa Fluor-488 (1:500, Abcam, ab150173), and Hoechst 33342 (1:5000, Invitrogen, H3570). Images of recorded neurons were obtained with a confocal microscope (Olympus Fluoview).

## RESULTS

### VPMpc afferents target GC neurons with laminar specific properties

We used an optogenetic approach to investigate the connectivity and synaptic properties of the projection from the gustatory thalamus, the parvicellular portion of the ventral posteromedial nucleus (VPMpc), to the gustatory cortex (GC). To determine the laminar distribution of VPMpc afferents to GC, we bilaterally injected CAG-AAV9-eYFP-ChR2 into the VPMpc of juvenile male and female rats. To characterize the synaptic properties of the VPMpc-GC input we performed whole-cell patch clamp recordings from excitatory neurons in different cortical layers and recorded excitatory postsynaptic currents (EPSCs) and potentials (EPSPs) evoked by stimulation of VPMpc axon terminals (Fig. 1). Recorded neurons were filled with biocytin for *post hoc* analysis of laminar location (see Methods; Fig. 1A, Cortical depth ratio, permutation t-test: Layer 4, n=73 vs Layer 2/3, n=11, *P*<1^-4^; vs Layer 5, n=28, *P*<1^-4^; vs Layer 6, n=17, *P*<1^-4^). In other sensory cortices, thalamocortical afferents primarily target neurons in Layer 4, and to a lesser extent, Layer 6. Similarly, in GC, visualization of the virally transfected VPMpc afferents showed densest terminal labeling in Layer 4 and Layer 6. However, VPMpc axons were also present in L2/3 and L5 and when we activated the VPMpc afferents, we found that optogenetic stimulation evoked EPSCs in neurons across all cortical layers (L2/3, L4, L5, L6, Fig. 1B, C). Overall, the percentage of neurons that received input from VPMpc did not differ between L4, L5, and L6, but a much smaller number of L2/3 neurons responded to VPMpc afferent stimulation (Fig. 1C, VPMpc connectivity %: Chi-square, χ^2^_(3)_=7.66, *P*= .05). The onset latencies of VPMpc-EPSCs onto neurons in all layers were indicative of a monosynaptic input yet demonstrated laminar specificity as VPMpc-EPSCs onto L4 neurons had a significantly shorter latency than those onto L5 or L6 neurons (Table 1, EPSC latency). The amplitude of VPMpc evoked responses onto neurons located in L4 was significantly larger compared to L2/3, L5, and L6 (Fig. 1C, Table 1). Consistent with the larger amplitude of L4 VPMpc-EPSCs, the total charge of the VPMpc-EPSCs onto L4 neurons was significantly larger than L2/3, L5, or L6 neurons, and the rise time of L4 currents was significantly shorter than those in L2 and L6 (Table 1, EPSC rise time). However, there were no laminar differences in the VPMpc-EPSC decay time constant (tau) (Table 1, EPSC charge, EPSC decay tau). The laminar-specific amplitude of VPMpc-EPSCs translated into laminar differences in the amplitude of evoked VPMpc-excitatory postsynaptic potentials (EPSPs), with larger changes in the membrane potential of L4 neurons compared to neurons in L2/3 and L5 (Fig. 1C, D, Table 1), indicating proportional input/output transformations. Thus, while thalamocortical input from VPMpc makes functional synapses with neurons in multiple GC layers, the synaptic properties differ based on neuronal location.

**Table 1:** Comparison of L4 VPMpc-EPSCs and VPMpc-EPSPs to L2, L5, and L6.

|  | **Layer 4** | **Layer 2** | **Layer 5** | **Layer 6** |
| --- | --- | --- | --- | --- |
| **EPSC amplitude (pA)**<br>*effect size vs. L4* | 311.81 ± 27.39<br>n=67 | 48.59 ± 11.6<br>-263.22 (-320.43, -205.76)<br>P=.004, n=7 | 106.05 ± 26.72<br>-205.76 (-270.45, -121.04)<br>P=.001, n=14 | 89.49 ± 33.53<br>-222.31 (-284.48, -98.27)<br>P=.006, n=8 |
| **EPSC charge (pA/ms)**<br>*effect size vs. L4* | 4.35 ± 0.37<br>n=66 | 0.93 ± 0.19<br>-3.42 (-4.24, -2.62)<br>P=.003, n=8 | 1.65 ± 0.37<br>-2.71 (-3.71, -1.62)<br>P=.001, n=14 | 1.54 ± 0.63<br>-2.81 (-3.79, -0.67)<br>P=.01, n=8 |
| **EPSC risetime (ms)**<br>*effect size vs. L4* | 3.75 ± 0.18<br>n=65 | 4.95 ± 0.77<br>1.20 (0.07, 3.02)<br>P=.05, n=7 | 4.51 ± 0.43<br>0.76 (-0.12, 1.64)<br>P=0.1, n=14 | 5.52 ± 0.84<br>1.76 (0.34, 3.51)<br>P=.008, n=8 |
| **EPSC latency (ms)**<br>*effect size vs. L4* | 2.37 ± 0.06<br>n=66 | 2.43 ± 0.20<br>0.06 (-0.34, 0.43)<br>P=.74, n=8 | 2.83 ± 0.21<br>0.46 (0.09, 0.91)<br>P=.005, n=14 | 3.16 ± 0.37<br>0.79 (0.20, 1.58)<br>P=.002, n=8 |
| **EPSC decay tau (ms)**<br>*effect size vs. L4* | 10.8 ± 0.4<br>n=67 | 10.4 ± 1.9<br>-0.4 (-3.7, 3.5)<br>P=.76, n=7 | 11.7 ± 1.0<br>0.9 (-1.2, 2.9)<br>P=.36, n=14 | 11.3 ± 1.5<br>0.5 (-1.7, 4.1)<br>P=.66, n=8 |
| **EPSP amplitude (mV)**<br>*effect size vs. L4* | 11.17 ± 0.89<br>n=61 | 3.63 ± 1.12<br>-6.08 (-8.21, -2.23)<br>P=.03, n=7 | 5.19 ± 0.88<br>-4.53 (-6.78, -1.91)<br>P=.03, n=13 | 6.51 ± 1.72<br>-3.21 (-6.29, 0.90)<br>P=.25, n=7 |
| **EPSC amp-animal (pA)**<br>*effect size vs. L4* | 302.03 ± 35.77<br>n=24 | 54.24 ± 7.20<br>-247.79 (-322.15, -183.67)<br>P=.003, n=5 | 110.33 ± 23.43<br>-191.70 (-272.11, -109.74)<br>P=.003, n=10 | 100.26 ± 43.89<br>-201.77 (-279.26, -59.04)<br>P=.01, n=6 |
| **EPSP amp-animal (mV)**<br>*effect size vs. L4* | 10.61 ± 1.24<br>n=23 | 4.82 ± 1.39<br>-5.79 (-8.53, -1.44)<br>P=.05, n=5 | 4.93 ± 0.99<br>-5.68 (-8.66, -2.64)<br>P=.008, n=10 | 6.50 ± 1.76<br>-4.11 (-8.02, -0.23)<br>P=.16, n=5 |
| **EPSC ppr 20hz, 40hz**<br>20hz vs 40hz | 1.11 ± 0.03, 0.60 ± 0.03<br>-0.51 (-0.57, -0.46)<br>P<.0001, n=56 | 1.16 ± 0.11, 0.71 ± 0.06<br>-0.45 (-0.69, -0.15)<br>P=.03, n=7 | 1.21 ± 0.06, 0.68 ± 0.08<br>-0.53 (-0.69, -0.40)<br>P<.0001, n=12 | 1.20 ± 0.11, 0.60 ± 0.15<br>-0.60 (-0.79, -0.42)<br>P=.01, n=7 |
| **EPSC ssr 20hz, 40hz**<br>20hz vs 40hz | 1.05 ± 0.04, 0.66 ± 0.04<br>-0.39 (-0.47, -0.32)<br>P<.0001, n=56 | 1.17 ± 0.12, 0.74 ± 0.04<br>-0.43 (-0.68, -0.24)<br>P=.03, n=7 | 1.18 ± 0.08, 0.77 ± 0.09<br>-0.41 (-0.50, -0.33)<br>P<.0001, n=12 | 1.45 ± 0.28, 0.83 ± 0.26<br>-0.62 (-0.93, -0.44)<br>P=.01, n=7 |

### VPMpc afferent input to GC is gated by distinct activity regimes

Previous studies reported that laminar differences in the strength of sensory thalamocortical input are accompanied by distinct short-term dynamics (Sherman and Guillery, 1998; Wang et al., 2013). Typically, thalamocortical input to L4 exhibits the strongest depression following repetitive stimulation, indicative of synaptic input that can reliably “drive” the circuit (Kloc and Maffei, 2014). To determine if similar properties apply to VPMpc-GC synapses, we recorded excitatory neurons while activating VPMpc axons with repetitive stimulation at 20Hz and 40Hz. Despite the laminar differences in VPMpc-EPSC amplitude, the short-term dynamics of VPMpc-EPSCs were comparable across layers, a unique feature of thalamocortical transmission in GC. There was a strong frequency dependence of short term dynamics that was consistent across neurons in all layers (Fig. 2A, 2-way ANOVA, frequency: F_(1,78)_=195.6, *P*<1^-4^; layer, F_(3,78)_=0.8229, *P*=.49; interaction, F_(3,78)_=0.4752, *P*=.7). Repetitive stimulation at 20Hz resulted in short-term facilitation of VPMpc-EPSCs in the majority of recorded neurons, whereas 40Hz stimulation resulted in short-term depression of VPMpc-EPSCs (facilitation/depression: 20Hz, 59/25; 40Hz, 8/76; Chi-square, χ^2^_(1)_ = 64.57, *P*<.0001). Activation of VPMpc afferents at 20Hz showed an increase (Fig. 2A, Table 1) in both paired-pulse ratio (PPR, epsc2/epsc1) and steady-state ratio (SSR, epsc5/epsc1), while stimulation at 40Hz produced a reliable decrease. This data suggests that the specific activity regime of VPMpc afferents can profoundly affect the recruitment of the GC circuit, and that the effect is uniform across cortical layers. To examine the frequency dependence of VPMpc afferent activation on GC neuronal output, we repetitively stimulated VPMpc axons while recording from GC neurons in current clamp. The trains of stimuli evoked a series of individual VPMpc-EPSPs that rode upon a slower, sustained membrane depolarization. When we quantified the VPMpc-EPSPs, we observed changes in PPR and SSR following the same trend as VPMpc-EPSCS at 40Hz and 20Hz (Fig. 2B) at synapses across all layers. The amplitude of the underlying depolarization also increased with each subsequent LED pulse and was significantly more pronounced at 40Hz for L2/3, L4, and L5 neurons (Fig. 2C, Table 1), indicating frequency dependent response summation. These results show that the frequency of incoming VPMpc inputs can gate the activation of the circuit in GC, as well as the magnitude of depolarization, or up state, in response to incoming activity.

**Figure 2:**
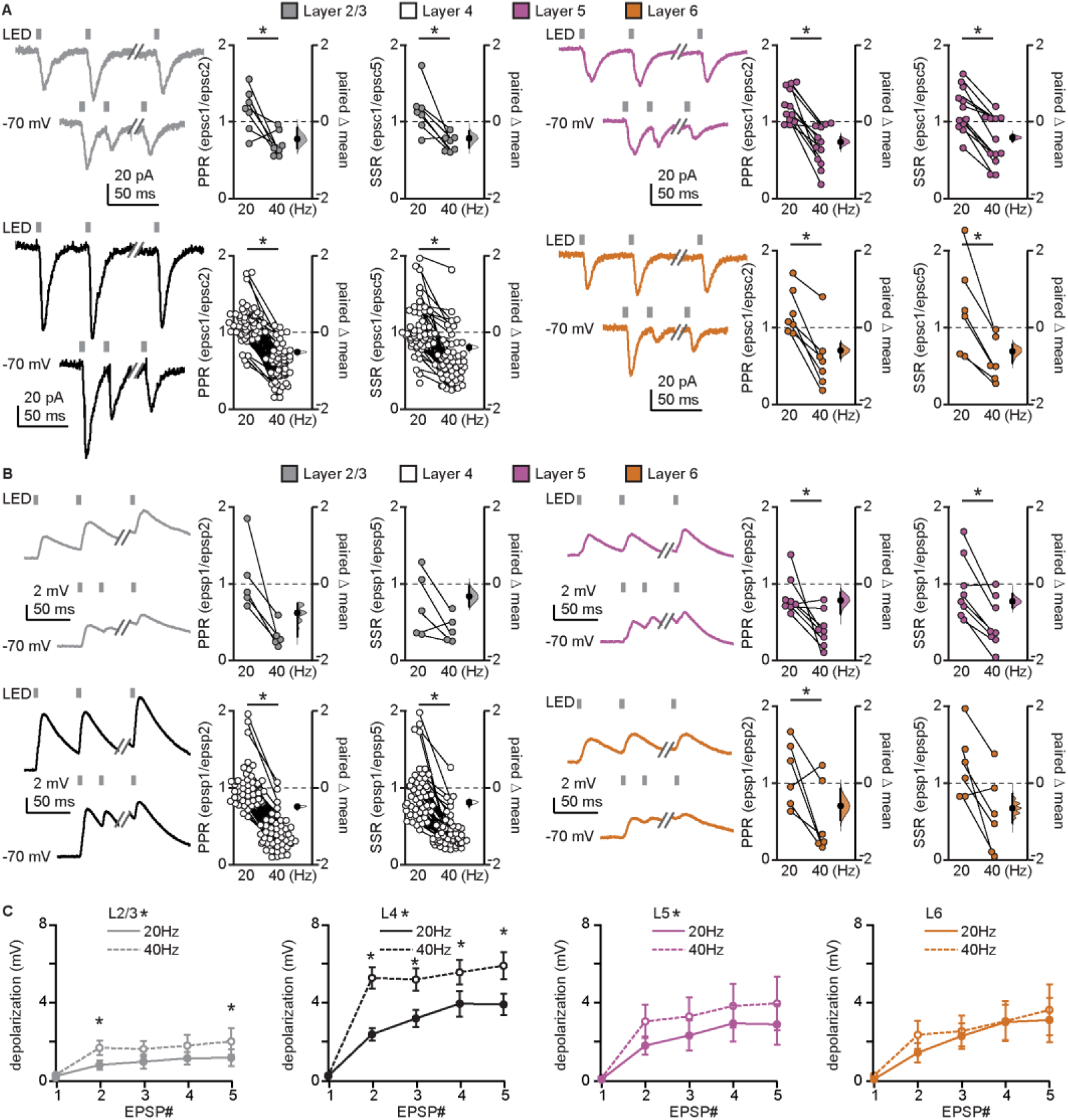
Activation frequency differentially gates VPMpc afferent input to GC. **A**. Top, left: Representative traces of VPMpc-EPSCs evoked following 20hHz (top) and 40Hz (bottom) repetitive stimulation of VPMpc afferents onto L2/3 neuron (grey, n=7) and summary of paired-pulse ratio (PPR, epsc2/epsc1) and steady-state ratio (SSR, epsc5/epsc1). Bottom, left: L4 neuron (black, n=56); Top, right: L5 neuron (magenta, n=12); Bottom, right: L6 neuron (orange, n=7). **B**. Top, left: Representative traces of VPMpc-EPSPs evoked following 20Hz (top) and 40Hz (bottom) repetitive stimulation of VPMpc afferents onto L2/3 neuron and summary of paired-pulse ratio (PPR, epsc2/epsc1) and steady-state ratio (SSR, epsc5/epsc1; n=5). Bottom, left: L4 neuron (n=50); Top, right: L5 neuron (n=8); Bottom, right: L6 neuron (n=6). **C**. Summary of tonic depolarization amplitude during VPMpc-EPSPs evoked at 20Hz (solid line) and 40Hz (dashed line). Grey: 2/3, n=5; black: L/4, n=50; magenta: L5, n=8; orange: L6, n=7. * indicates p ≥ 0.05. Error bars ± SEM.

### VPMpc afferents target specific subclasses of L4 GABAergic neurons

A factor that can gate how long-range inputs recruit local cortical circuits is GABAergic inhibition. Sensory thalamocortical afferents in other areas strongly innervate fast-spiking, parvalbumin-expressing GABAergic neurons (FS) (Cruikshank et al., 2007; Kloc and Maffei, 2014) and higher-order thalamic nuclei can also innervate regular-spiking non-pyramidal GABAergic neurons (RSNP) (Tan et al., 2008; Cruikshank et al., 2012; Audette et al., 2018). We recorded from both FS and RSNP neurons to determine if VPMpc targets one, or both, of these groups in L4 of GC. Identity of the recorded neurons was determined online by assessing the firing pattern during steady-state current injections, and later confirmed via post-hoc morphological analysis and immunostaining for GAD-67 (Fig. 3A). Optogenetic stimulation of VPMpc afferents elicited EPSCs and EPSPs in 35% of FS neurons, a smaller proportion compared to PYR neurons (Fig. 3B, C, VPMpc connectivity %: Chi-square, χ^2^_(1)_=14.95, *P*=.0001), while no responses were detected in RSNP neurons. The onset latency of VPMpc-EPSCs onto FS neurons was consistent with a monosynaptic input and comparable to the onset latency for PYR neurons (Fig. 3C, EPSC onset latency: permutation t-test, *P*=.98). Likewise, the amplitude of VPMpc-EPSCs and VPMpc-EPSPs did not differ between the PYR and FS groups (Fig. 3C, EPSC amplitude: permutation t-test, *P*=.09; EPSP amplitude: permutation t-test, *P*=.23), indicating proportional input/output transformations in both neuron groups. The total charge of VPMpc-EPSCs, however, did differ between FS and PYR neurons, as VPMpc-EPSCs onto FS neurons had a significantly faster decay time constant resulting in a smaller overall charge despite the comparable amplitude (Fig. 3C, EPSC charge: permutation t-test, *P*=.05; EPSC decay tau: permutation t-test, *P*=.0002). These results indicate that VPMpc afferents directly target GABAergic FS neurons in GC, and the evoked responses exhibit similar amplitudes, but faster kinetics and reduced charge transfer compared to VPMpc inputs onto excitatory neurons.

**Figure 3:**
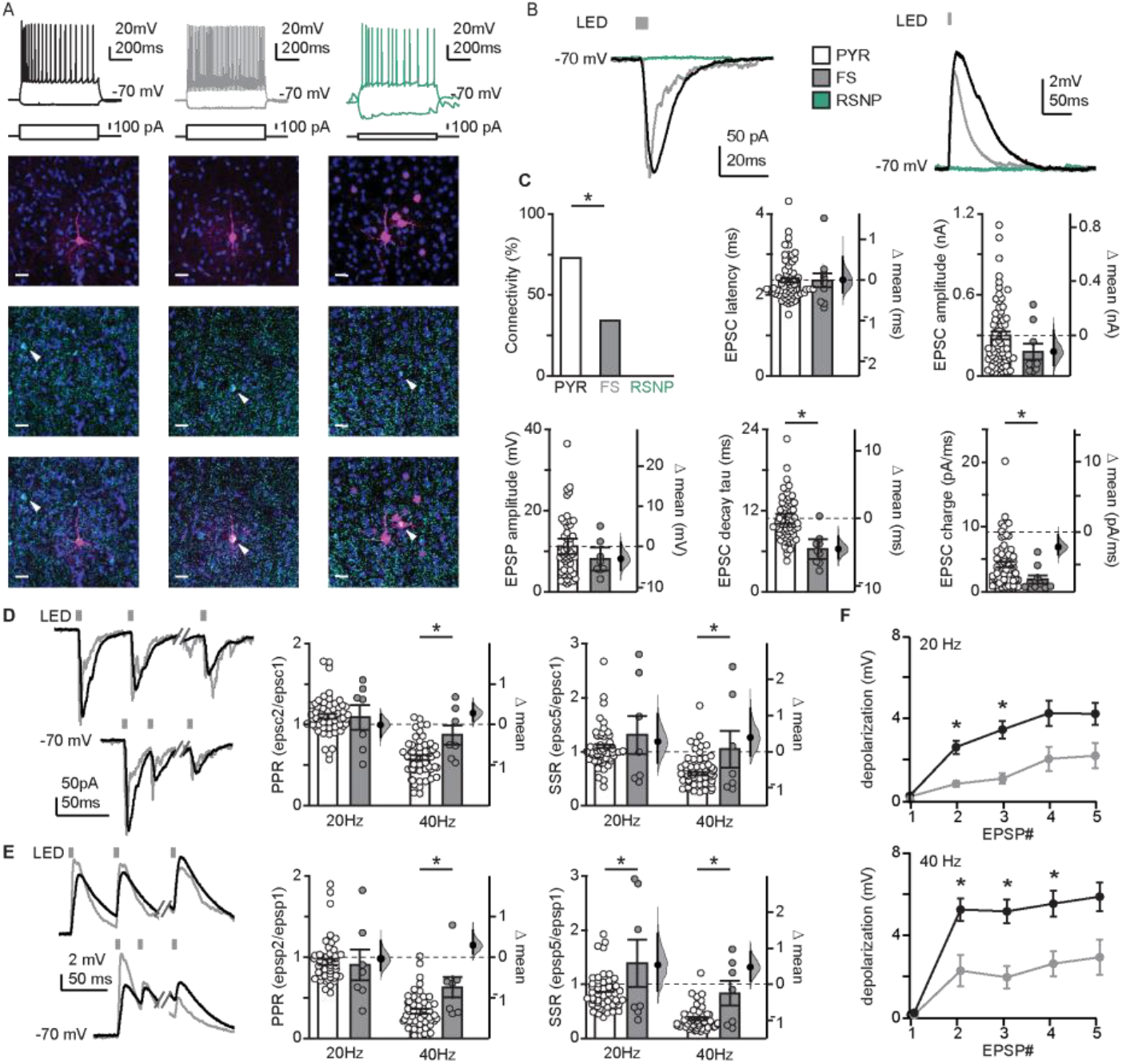
VPMpc afferents recruit GABAergic neurons in GC with cell-type specific dynamics. **A**. Left: sample traces of firing pattern of L4 PYR to steady-state current injection (black) and image of a biocytin-filled L4 PYR (note lack of co-localization with GAD-67). Middle: sample traces of firing pattern of L4 FS neuron to steady-state current injection (grey) and image of a biocytin-filled L4 FS (co-labeled by GAD-67). Right: sample traces of firing pattern of L4 RSNP neuron to steady-state current injection (green) and image of a biocytin-filled L4 RSNP neuron co-labeled with GAD-67 (blue: Hoechst, red: biocytin, green: GAD-67. Scale bar = 20μm). **B**. Top: Sample traces of VPMpc-EPSCs (left) and VPMpc-EPSPs (right) onto L4 PYR (black), FS neuron (gray), and RSNP neuron (green) recorded at -70 mV. **C**. Top: Summary of VPMpc connection probability onto L4 PYR (n= 87/119), and FS (n=11/32) and RSNP (n=0/8) neurons; summary of VPMpc-EPSC onset latency and amplitude onto L4 PYR (n=66) and FS neurons (n=9). Bottom: Summary of VPMpc-EPSP amplitude (right) onto L4 PYR (n=52) and FS neurons (n=8); summary of VPMpc-EPSC charge and decay tau onto L4 PYR (n=66) and FS neurons (n=9); **D**. Left: Sample traces of VPMpc-EPSCs onto L4 PYR (black) and FS neurons (grey) evoked by repetitive stimulation of VPMpc afferents at 20Hz (top) and 40Hz (bottom). Right: Summary of paired-pulse ratio (PPR, epsc2/epsc1) and steady-state ratio (SSR, epsc5/epsc1; PYR, n=56; FS, n=7). **E**. Left: Sample traces of VPMpc-EPSPs onto L4 PYR (black) and FS inhibitory (grey) evoked by repetitive stimulation of VPMpc afferents at 20Hz (top) and 40Hz (bottom). Right: Summary of paired-pulse ratio (PPR, epsp2/epsp1) and steady-state ratio (SSR, epsp5/epsp1; PYR, n=49; FS, n=7). **F**. Summary of tonic depolarization amplitude during VPMpc-EPSPs evoked at 20Hz (top) and 40Hz (bottom) onto L4 PYR (black, n=49) and FS neurons (grey, n=7) recorded at -70 mV. * indicates p ≤ 0.05. Error bars ± SEM.

Cell-type specific short-term dynamics of thalamocortical circuits can modulate cortical activation in response to sensory stimuli. When we examined the short-term dynamics of VPMpc-EPSCs onto inhibitory FS neurons, there was a frequency-dependent decrease of both PPR and SSR at 40Hz compared to 20Hz, similar to what was observed in PYR neurons (Fig. 3D, 2-way ANOVA, PPR: frequency: F_(1,61)_=71.01, *P*<1^-4^; cell-type, F_(1,61)_=2.483, *P*=.12; interaction, F_(1,61)_=11.41, *P*=.0013; post hoc Sidak’s, 20Hz vs 40Hz, PYR *P*<1^-4^, FS *P*=.02; SSR: frequency: F_(1,61)_=78.42, *P*<1^-4^; cell-type, F_(1,61)_=5.941, *P*=.02; interaction, F_(1,61)_=7.740, *P*=.007; post hoc Sidak’s, 20Hz vs 40Hz, PYR *P*<.0001, FS *P*=.004). When we directly compared FS PPR and SSR to the PYR population, we found comparable values at 20Hz (Fig. 3D, post hoc Sidak’s, PYR vs FS, 20Hz PPR, *P*=.9; 20Hz SSR *P*=.3). However, at 40Hz, VPMpc synapses onto FS neurons showed significantly less depression compared to PYR (Fig. 3D, post hoc Sidak’s, PYR vs FS, 40Hz PPR, *P*=.009; 40Hz SSR *P*=.004).

Analysis of short-term dynamics of evoked voltage responses showed a pattern comparable to that described for currents, indicating proportional input/output transformations. VPMpc-EPSPs in FS neurons showed frequency-dependent changes in PPR and SSR, indicating increased short term depression between 20Hz and 40Hz (Fig. 3E, 2-way ANOVA, PPR: frequency: F_(1,54)_=62.37, *P*<1^-4^; cell-type, F_(1,54)_=1.889, *P*=.2; interaction, F_(1,54)_=8.938, *P*=.004); post hoc Sidak’s, 20Hz vs 40Hz, PYR *P*<1^-4^, FS *P*=.02; SSR: frequency: F_(1,55)_=36.77, *P*<1^-4^; cell-type, F_(1,55)_=13.26, *P*=.0006; interaction, F_(1,55)_=0.1107, *P*=.7; post hoc Sidak’s, 20Hz vs 40Hz, PYR *P*<1^-4^, FS *P*=.002). When we compared VPMpc-EPSPs dynamics between FS neurons and PYR neurons, there was no difference in PPR at 20Hz, however the 20Hz SSR and both PPR and SSR at 40Hz were larger in FS neurons, indicating reduced short-term depression of VPMpc synaptic inputs to FS neurons (Fig. 3E, posthoc Sidak’s, PYR vs FS, 20Hz PPR, *P*=.9; 20Hz SSR *P*=.003; 40Hz PPR, *P*=.02; 40Hz SSR *P*=.009). We also observed that the distinct kinetics of VPMpc-EPSPs onto FS and PYR neurons resulted in different degrees of voltage summation evoked by trains of stimuli: the amplitude of the underlying voltage depolarization for FS neurons was smaller compared to PYR neurons, and showed no difference across stimulation frequencies (Fig. 3F, 2-way ANOVA, frequency: F_(3,550)_=1.171, *P*<1^-4^; EPSP#, F_(4,550)_=9.229, *P*<1^-4^; interaction, F_(12,550)_=1.171, *P*=.3); posthoc Tukey’s, 20Hz vs 40Hz, PYR *P*<1^-4^, FS *P*=.8; PYR vs FS, 20Hz *P*=.03, 40Hz *P*=.004). These distinctions in cell-type specific dynamics endow VPMpc-GC circuits with a high sensitivity to incoming patterned activity.

These results differ significantly from those reported for visual, auditory, and somatosensory cortices, where thalamocortical inputs onto FS neurons are typically more powerful and show more short-term depression than those onto PYR neurons (Reichova and Sherman, 2004; Gabernet et al., 2005; Kloc and Maffei, 2014), highlighting important differences in thalamocortical activation of the gustatory insular cortex compared to other sensory areas.

### VPMpc afferents recruit strong feedforward inhibition in GC

As VPMpc afferents recruit FS neurons, we set out to assess the extent of VPMpc-evoked feedforward inhibition onto GC L4 PYR neurons. To isolate excitatory and inhibitory postsynaptic currents from PYR neurons we used an internal solution that allowed us to obtain stable recordings at the reversal potential for chloride (−50mV in our experimental conditions) and at the reversal potential for excitatory responses (+10mV; Fig. 4A). In all cells, when recording at +10mV, we observed a large, disynaptic IPSC in response to optogenetic stimulation of VPMpc-afferents. VPMpc-IPSCs had a significantly longer onset latency and larger amplitude than the monosynaptic VPMpc-EPSCs (Fig. 4A, PSC latency: paired permutation t-test, EPSC vs. IPSC, *P*=.0004; PSC amplitude: paired permutation t-test, EPSC vs. IPSC, *P*=.0004). We then quantified the evoked E/I ratio, which was dominated by inhibitory drive (Fig. 4A, E/I ratio = 0.3 ± 0.04, n = 11). These data suggest that disynaptic inhibition evoked by VPMpc may shunt the excitatory VPMpc postsynaptic response. We compared short-term dynamics of VPMpc-evoked excitation and feedforward inhibition to examine if the E/I ratio changes during repetitive stimulation. While the E/I ratio of evoked responses during 20Hz and 40Hz stimuli trains were dominated by inhibition, frequency-dependent differences emerged towards the end of train where 20Hz stimulation resulted in a larger E/I ratio than 40Hz stimulation (Fig. 4B, 2-way ANOVA, frequency: F(1,31)=15.72, P=.0004; PSC, F(4,31)=1.612, P=.2; interaction, F(4,31)=5.828, P=.1). When we examined the PPR and SSR, we observed a large decrease in VPMpc-EPSC PPR from 20Hz to 40Hz, and a more modest decrease in the IPSC PPR (Fig.4B, 2-way ANOVA, PPR, frequency: F_(1,11)_=35.05, *P*=1^-4^; PSC, F_(1,11)_=2.043, *P*=.2; interaction, F_(1,11)_=5.828, *P*=.03; *post hoc* Sidak’s, 20Hz vs 40Hz, EPSC P=.0003, IPSC P=.05). The decrease in SSR for VPMpc-EPSC at 20Hz and 40Hz followed the same trend as PPR. Differently, the SSR for IPSCs was not affected by the frequency of stimulation and was comparable at 20Hz and 40Hz (Fig. 4B, 2-way ANOVA, SSR, frequency: F_(1,13)_=54.55, *P*<1^-4^; PSC, F_(1,13)_=2.723, *P*=.12; interaction, F_(1,13)_=21.89, *P*=.0004; posthoc Sidak’s, 20Hz vs 40Hz, EPSC *P*<1^-4^, IPSC *P*=.13). The VPMpc-EPSC PPR and SSR at 20Hz were larger than the 20Hz PPR and SSR for inhibition (Fig 4B, posthoc Sidak’s, VPMpc-EPSC vs IPSC, PPR 20Hz *P*=.04, 40Hz *P*=.9; SSR 20Hz *P*=.01, 40Hz *P*=.9). Activation of VPMpc inputs in GC, thus, recruits excitatory and inhibitory feedforward circuits evoking postsynaptic responses with distinct time constants and short-term dynamics. The brief di-synaptic delay for the activation of feedforward inhibition is consistent with previous findings reporting an early contribution of inhibition to the raising phase of the evoked postsynaptic potential recorded *in vivo* (Stone et al., 2020). The differences in decay time constant and short term dynamics of feedforward excitation and inhibition suggest that repetitive activation of VPMpc afferents by patterns of stimuli can delineate precise temporal windows for the integration of incoming inputs in PYR neurons (Wehr and Zador, 2003).

**Figure 4:**
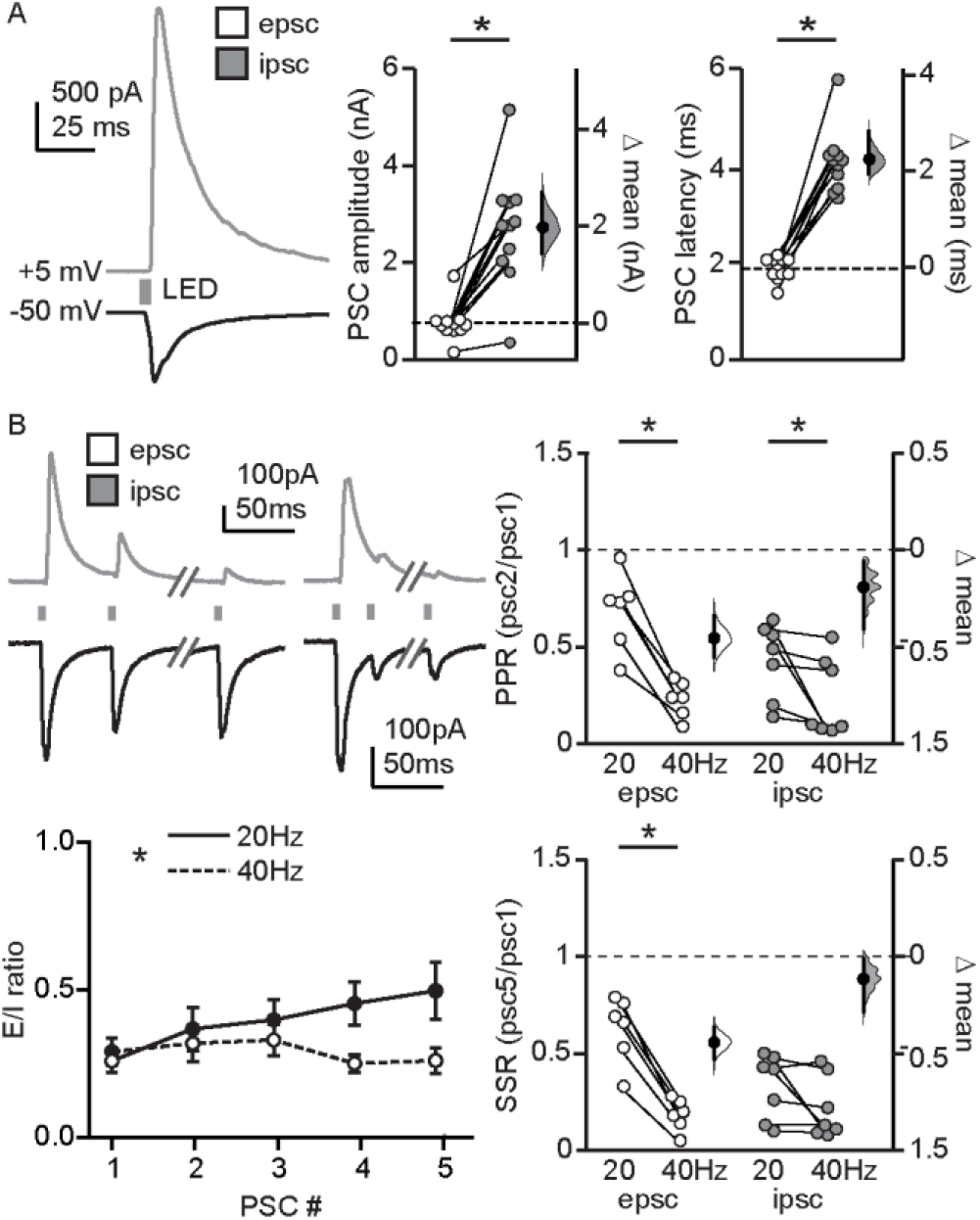
VPMpc afferent activation drives feedforward inhibition onto L4 excitatory neurons in GC. **A**. Left: Representative traces of VPMpc-IPSCs (grey) and VPMpc-EPSCs (black) onto a L4 excitatory neuron. Right: Summary of amplitude and onset latency of VPMpc-EPSCs (white) and IPSCs (grey) onto L4 excitatory neurons (n=8). **B**. Left, top: Sample traces of VPMpc-IPSCs (grey) and VPMpc-EPSCs (black) onto L4 excitatory neuron evoked following repetitive stimulation of VPMpc afferents at 20Hz (left) and 40Hz (right). Left, bottom: Summary of E/I ratio (n=8) following repetitive stimulation of VPMpc afferents at 20Hz (solid line) and 40Hz (dashed line). Right: Summary of paired-pulse ratio (top; PPR, psc2/psc1) and steady-state ratio (bottom; SSR, psc5/psc1) at 20Hz and 40Hz for VPMpc-EPSCs (white, n=8) and IPSCs (grey, n=8). * indicates p ≤ 0.05. Error bars ± SEM.

### Inhibitory synaptic transmission gates VPMpc input to GC

VPMpc afferents in GC directly recruit GABAergic FS neurons and drive strong feedforward inhibition onto PYR neurons, indicating that VPMpc activity engages GC inhibitory circuits. To further investigate how cortical inhibition may modulate thalamocortical inputs to GC, we used GABA receptor agonists to test the effects of increased cortical inhibition on VPMpc-EPSCs (Fig. 5A, B). GABAergic transmission is mediated by fast-acting, ionotropic GABA_A_ receptors and slower, metabotropic GABA_B_ receptors. To broadly activate GABA_A_ receptors we bath applied muscimol, while baclofen was used to activate GABA_B_ receptors. Bath application of muscimol significantly reduced the amplitude of the VPMpc-EPSCs (Fig. 5A, EPSC amplitude: paired permutation t-test, *P*=.002). The reduction in VPMpc amplitude was accompanied by a shift in the holding current, as well as a significant decrease in the decay tau of the VPMpc-EPSC indicating that GABA_A_ receptors are present postsynaptically (Fig. 5A, holding current: paired permutation t-test, *P*=.02; decay tau: paired permutation t-test, *P*=1^-4^). To determine if the effect of muscimol on amplitude was fully accounted for by the shift in holding current, we measured the correlation between change in amplitude and change in holding current. There was no correlation between these parameters (Fig. 5C, Δ EPSC amplitude vs Δ holding current: Spearman rank order correlation, muscimol, *P*=.25), indicating that an additional mechanism was recruited by activation of GABA_A_ receptors with muscimol. Next, we tested the effect of activation of GABA_B_ receptors on VPMpc-EPSCs by bath application of baclofen. Baclofen also induced a significant reduction in the amplitude of VPMpc-EPSCs (Fig. 5B, EPSC amplitude: paired permutation t-test, *P*=.005), as well as a shift in the holding current (Fig. 5B. holding current: paired permutation t-test, *P*=.82). However, differently from muscimol, bath application of baclofen had no effect on the VPMpc-EPSC decay tau (Fig. 5B, decay tau: paired permutation t-test, *P*=.61). In the case of baclofen, we also assessed whether the change in holding current could account for the decrease in VPMpc-EPSC amplitude. As for muscimol, there was no correlation between the change in holding current and change in amplitude, suggesting an additional site for GABAergic modulation of VPMpc-EPSC amplitude (Fig. 5C, Δ EPSC amplitude vs Δ holding current: Spearman rank order correlation, baclofen, *P*=.18).

**Figure 5:**
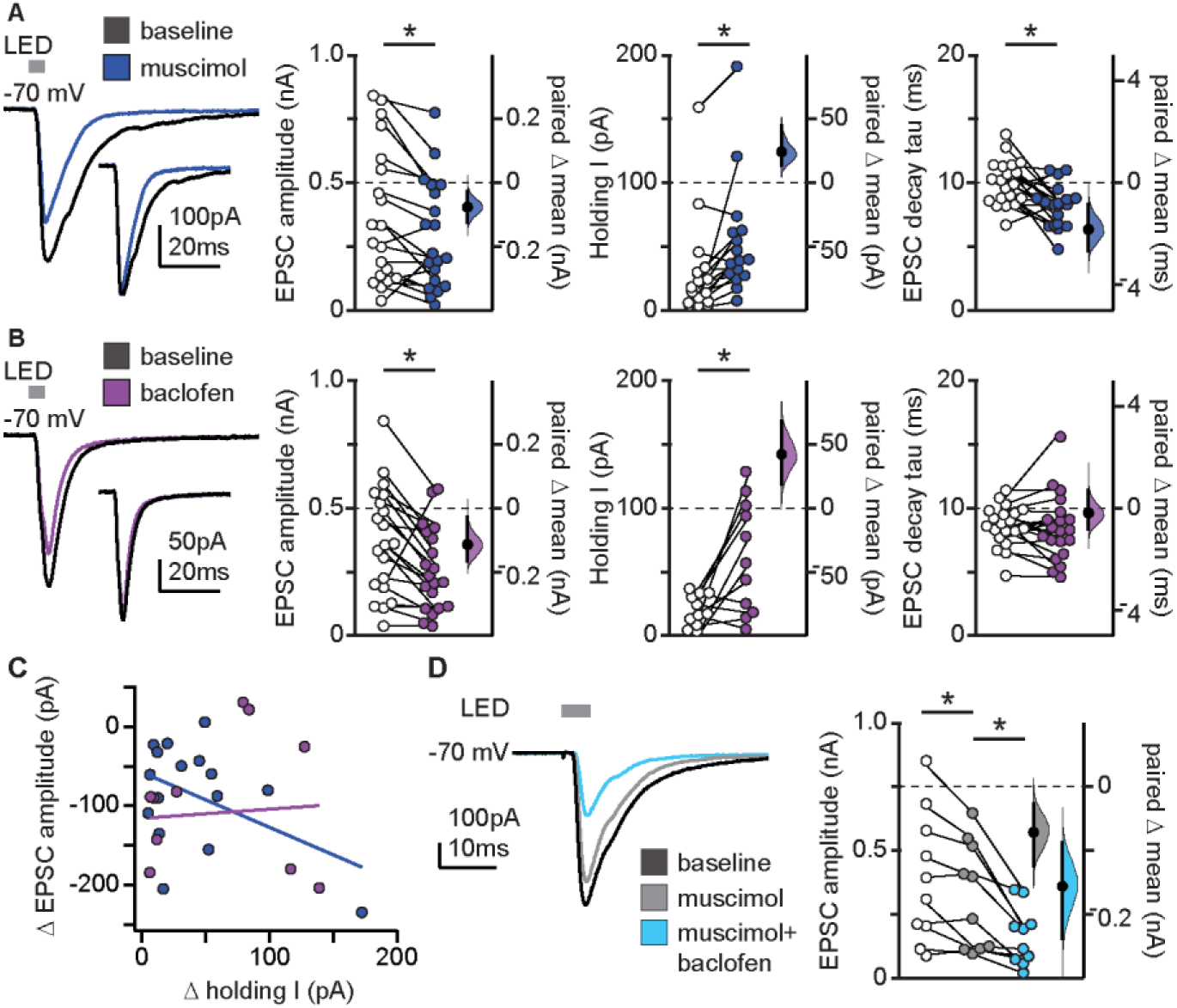
GABA gates VPMpc input in GC. **A**. Left: Sample traces of VPMpc-EPSCs onto L4 pyramidal neuron at baseline (black) and during bath application of muscimol (blue, n=20). Inset: VPMpc-EPSCs recorded with muscimol trace scaled to baseline to compare decay kinetics. Right: Summary of VPMpc-EPSC amplitude, holding current, and decay tau in muscimol. **B**. Left: Sample traces of VPMpc-EPSCs onto L4 pyramidal neuron at baseline (black) and during bath application of baclofen (purple, n=20). Insets: VPMpc-EPSCs recorded with baclofen trace scaled to baseline to compare decay kinetics. Right: Summary of VPMpc-EPSC amplitude, holding current, and decay tau in baclofen. **C**. Plot of the change in VPMpc-EPSC amplitude versus shift in holding current following drug application (blue: muscimol, purple: baclofen). **D**. Left: Sample traces of VPMpc-EPSCs following consecutive and additive application of muscimol (grey) followed by muscimol and baclofen (light blue). Right: Summary of VPMpc-EPSC amplitude in the presence of muscimol (grey) and muscimol and baclofen combined (light blue, n=11). * indicates p ≤ 0.05. Error bars ± SEM.

To ensure that the actions of muscimol and baclofen did not result from nonspecific receptor cross-activation, in a subset of experiments we sequentially and additively bath applied muscimol and baclofen (Fig. 5D). We observed a decrease in VPMpc-EPSC amplitude following bath application of muscimol and an additional decrease in VPMpc-EPSC amplitude when muscimol and baclofen were co-applied, indicating that both muscimol and baclofen have distinct and additive actions on VPMpc-EPSCs in GC (Fig. 5D, EPSC amplitude: paired permutation t-test, muscimol + baclofen, *P*<1^-4^). This effect is comparable to that we previously reported in rodent primary visual cortex (Wang et al., 2019), and points to the presence of presynaptic GABA_A_ receptors on thalamocortical terminals.

### Cortical GABAergic feedback presynaptically gates VPMpc afferents in GC

Whereas presynaptic GABA_B_ receptors are common in cortex, there are fewer reports of presynaptic GABA_A_ receptors (Ruiz et al., 2010; Wang et al., 2019). To determine if a presynaptic modulation by inhibition could account for the decrease in VPMpc-EPSC amplitude in response to the application of GABA receptor agonists, we recorded from PYR neurons using repetitive stimulation of VPMpc afferents before and after bath application of muscimol or baclofen (Fig. 6A, B). We assessed the PPR and SSR on trains of VPMpc-EPSCs recorded before and after drug delivery. There were significant changes in both PPR and SSR following application of either muscimol or baclofen, pointing to changes in release probability and supporting the presence of a presynaptic site for GABA action (Fig. 6A, B). Application of muscimol consistently decreased PPR and SSR both at 20Hz and 40Hz (Fig. 6A, paired permutation t-test, baseline vs. muscimol PPR: 20Hz, *P*=.002; 40Hz, *P*=.0004; SSR: 20Hz, *P*<1^-4^; 40Hz, *P*<1^-4^), whereas baclofen application resulted in an increase in PPR and SSR at both frequencies (Fig. 6B, paired permutation t-test, baseline vs. baclofen PPR: 20Hz, *P*=.009; 40Hz, *P*=.001; SSR: 20Hz, *P*<1^-4^; 40Hz, P=.0004). Because presynaptic activation of GABA_A_ and GABA_B_ receptors produced opposite changes in short-term dynamics of VPMpc-EPSCs, these presynaptic effects were almost completely occluded when the drugs were co-applied (Fig. 6C). Co-application of muscimol and baclofen had no effect on the PPR at 20Hz or 40Hz, or the SSR at 40Hz, but resulted in a net increase in the SSR at 20Hz (Fig. 6C, paired permutation t-test, baseline vs. muscimol + baclofen PPR: 20Hz, *P*=0.7; 40Hz, *P*=.08; SSR: 20Hz, *P*=.005; 40Hz, *P*=.06). These results indicate that the presence of presynaptic GABA_A_ and GABA_B_ receptors on VPMpc afferent terminals provide a distinct mechanism for control of VPMpc input to GC via inhibitory cortical feedback.

**Figure 6:**
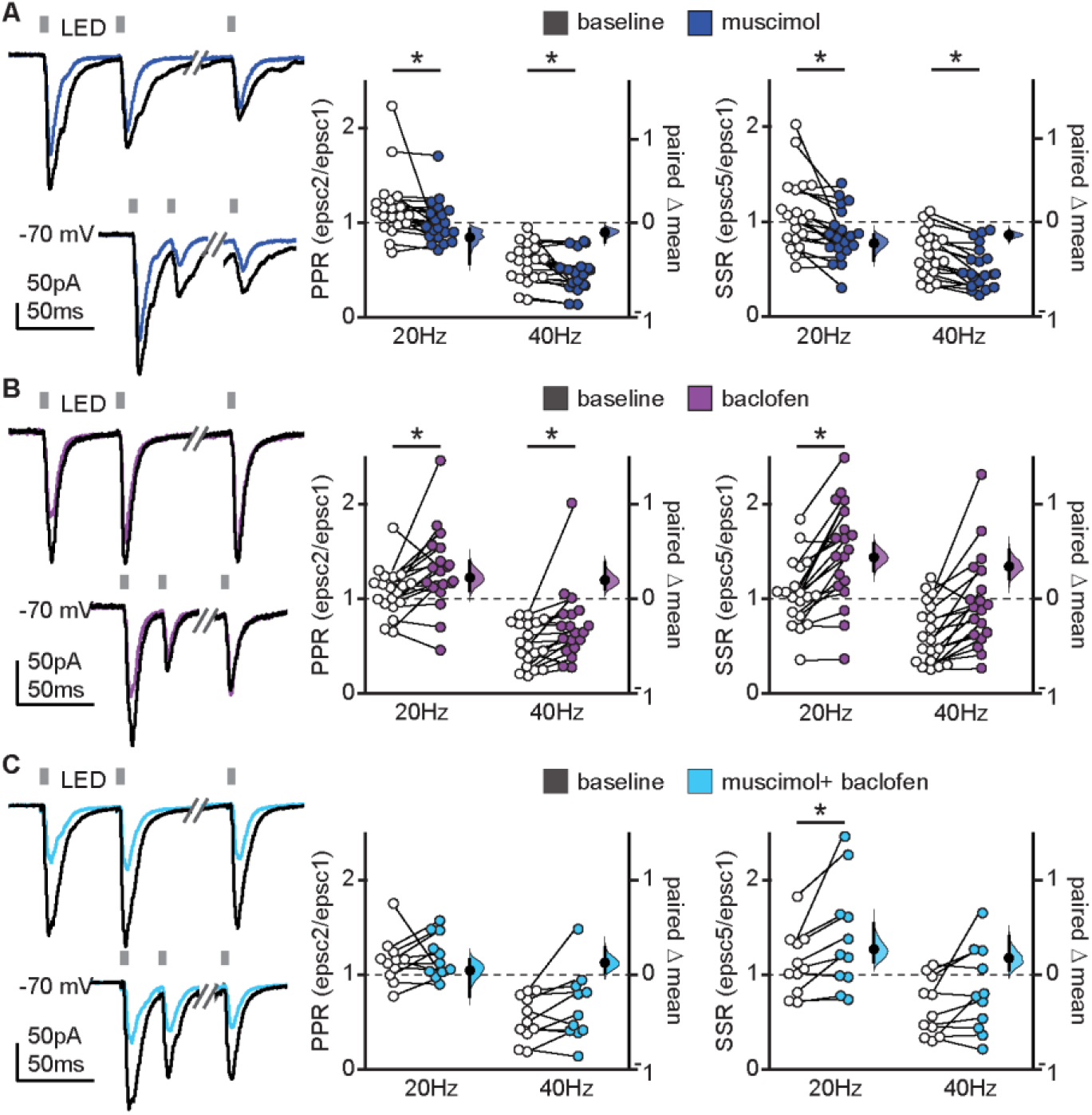
Cortical GABAergic feedback presynaptically gates VPMpc afferents in GC. **A**. Left: Sample traces of VPMpc-EPSCs evoked following repetitive stimulation of VPMpc afferents onto L4 pyramidal neuron at baseline (black) and during bath application of muscimol (blue, n=20). Right: Summary of paired-pulse ratio (PPR, epsc2/epsc1) and steady-state ratio (SSR, epsc5/epsc1) at baseline (black) and in the presence of muscimol (blue). **B**. Left: Sample traces of VPMpc-EPSCs evoked following repetitive stimulation of VPMpc afferents onto L4 pyramidal neuron at baseline (black) and during bath application of baclofen (purple, n=19). Right: Summary of paired-pulse ratio (PPR, epsc2/epsc1) and steady-state ratio (SSR, epsc5/epsc1) at baseline (black) and in the presence of baclofen (purple). **C**. Left: Sample traces of VPMpc-EPSCs evoked following repetitive stimulation of VPMpc afferents onto L4 pyramidal neuron at baseline (black) and during simultaneous bath application of muscimol and baclofen (light blue, n=11). Right: Summary of paired-pulse ratio (PPR, epsc2/epsc1) and steady-state ratio (SSR, epsc5/epsc1) at baseline (black) and in the presence of muscimol and baclofen (light blue). * indicates p ≥ 0.05. Error bars ± SEM.

## DISCUSSION

In this study we investigated the synaptic organization and properties of thalamocortical projections from VPMpc to GC. Functional VPMpc-GC synapses were identified spanning cortical layers 2-6, with the most powerful input onto excitatory neurons in Layer 4. The L4 VPMpc-GC synaptic responses showed frequency-dependent short-term plasticity and were strongly gated by both feedforward and feedback inhibition. These results indicate that distinct patterns of thalamic activity can recruit both excitatory and inhibitory circuits in GC. Furthermore, our data provide insight into possible mechanisms employed by GC circuits to multiplex taste-related information and integrate diverse inputs.

### VPMpc modulation of GC network state

Previous studies have shown that inactivating VPMpc has a profound impact on the GC network state, dramatically altering the spectral content of cortical LFPs. Specifically, silencing VPMpc decreases the power in the gamma range, increases the power of lower frequency oscillations and has heterogeneous effects on the rate of spontaneous firing of GC neurons. These effects are reminiscent of sleep-related and anesthesia-related rhythms resulting from thalamic hyperpolarization (Samuelsen et al., 2013). Oscillatory rhythms can gate information transfer in cortical circuits, and can be generated via long-range inputs, local recurrent intracortical circuits, or a combination of both. Our data identify multiple local cortical mechanisms by which VPMpc afferents can modulate the GC network state. Firstly, we provide evidence that VPMpc afferents directly recruit FS GABAergic neurons in GC. Activity of cortical FS GABAergic neurons contributes to the generation of gamma oscillations (Cardin et al., 2009), thus their recruitment by thalamocortical axons is one way in which VPMpc input can contribute to fast, desynchronized oscillatory activity in GC. Conversely, quenching of this mechanism can explain the reduction of gamma oscillations seen after thalamic inactivation (Samuelsen et al., 2013). Secondly, our results show an effect of muscimol and baclofen on VPMpc-EPSCs, unveiling a contribution of GABA_A_ and GABA_B_ receptors to thalamocortical interactions in GC via postsynaptic modulation of GC neurons and presynaptic modulation of VPMpc axon terminals. Fast and slow GABAergic transmission via GABA_A_ and GABA_B_ receptors can trigger transitions between up and down states and can influence cortical synchrony (Mann et al., 2009; Sanchez-Vives et al., 2021). Thus, VPMpc recruitment of both feedforward thalamocortical and feedback corticothalamic inhibition via GABA_A_ and GABA_B_ receptors represent intra-GC mechanisms by which VPMpc inputs can influence the GC state.

### Temporal dynamics of VPMpc-GC synapses

The specific activity regime of thalamocortical neurons can also drive changes in the state of GC. While details of the firing patterns and rates of VPMpc neurons are unclear, there is some evidence that *in vivo* VPMpc neurons respond to tastant delivery with firing rates across a range of frequencies (Verhagen et al., 2003; Liu and Fontanini, 2015). We chose two frequencies within this range, 20Hz and 40Hz, and used trains of stimuli to investigate the frequency dependence of VPMpc-GC synaptic responses. We identified a pronounced effect of stimulation frequency on VPMpc-GC short-term synaptic dynamics. Following repetitive stimulation of VPMpc afferents in GC, quantification of the PPR and SSR revealed synaptic facilitation at 20Hz and depression at 40Hz. In other sensory systems, thalamocortical synapses from primary sensory thalamic afferents strongly drive the cortical circuits typically showing synaptic depression at most frequencies (Sherman and Guillery, 1998; Kloc and Maffei, 2014). This effect has been interpreted as depending on a high probability of neurotransmitter release at the first stimulation followed by progressive depletion of the availability of releasable vesicles with subsequent stimuli in the train. Short term facilitation, on the other hand, is often associated with modulatory thalamocortical afferents from secondary thalamic nuclei (Sherman and Guillery, 1998). The frequency dependence of thalamocortical short term dynamics – with one frequency inducing facilitation and another depression – has not been previously reported in thalamocortical systems and may be a unique property of chemosensory thalamocortical circuits. Based on these findings, it appears that VPMpc may function both as a driving and modulating thalamic nucleus and engage GC with distinct dynamics depending on incoming activity patterns. An alternative possibility is that the facilitating synaptic dynamics we observed at 20Hz are not necessarily a property of the VPMpc-GC synapse, but rather an epiphenomenon arising from external factors. Indeed, repetitive stimulation in experiments using a cesium internal solution to isolate the excitatory and inhibitory components of the VPMpc-EPSC showed synaptic depression at 20Hz. This observation suggests that the facilitating response observed at resting membrane potential may be due to modulation of the short-term plasticity of VPMpc evoked excitatory responses by cortical inhibition (Cruikshank et al., 2007; Yamamoto et al., 2010). This interpretation is further supported by the presence of GABA_A_ and GABA_B_ receptors on presynaptic VPMpc terminals, whose activation can directly modulate glutamate release properties at thalamocortical synapses.

### Temporal filtering and integration of diverse inputs

Feedforward inhibition can serve to gate the recruitment of cortical circuits by delimiting temporal windows for the activation of postsynaptic neurons (Gabernet et al., 2005). Therefore, while VPMpc-evoked feedforward inhibition acts to temporally filter VPMpc-evoked excitation, it can also affect the integration of other inputs onto the neuron. For example, VPMpc feedforward inhibition can increase the threshold of activation of the postsynaptic neuron, increasing the signal to noise ratio of the synaptic inputs that follow. Corticothalamic feedback inhibition, on the other hand, relies on presynaptic GABA_A_ and GABA_B_ receptors on VPMpc afferent terminals, dampening release (Wang et al., 2019). While the recruitment of inhibition by VPMpc afferents can generate the GABA release necessary to drive corticothalamic feedback, other inputs to GC have also been shown to recruit inhibitory circuits. For example, input from amygdalocortical synapses in GC, in addition to targeting pyramidal neurons, target both PV-expressing and SOM-expressing GABAergic neurons, recruiting feedforward inhibition (Haley et al., 2016). If thalamocortical evoked responses are preceded by amygdalocortical input, the amygdala-dependent increase in cortical GABAergic tone could gate VPMpc-GC synaptic activity and modulate thalamocortical inputs onto GC neurons. Indeed, work that examined the integration of BLA and VPMpc input onto GC neurons *in vivo* reported that repetitive activation of BLA afferents could modulate VPMpc evoked responses onto GC pyramidal neurons, leading to facilitation or suppression of the VPMpc-PSP depending on the magnitude of the inhibitory component of evoked responses (Stone et al., 2020). The results we report in this study provide further evidence of the importance of GC inhibitory circuits for the gating of sensory information as well as for the modulation of the integration of thalamocortical response dynamics.

### Comparison with other thalamocortical systems

When examined in the context of other thalamocortical circuits, VPMpc-GC inputs show similarity in the postsynaptic targets but display fundamental differences in synaptic organization. In neocortical sensory circuits, primary thalamocortical afferents are densely distributed in L4 and to a lesser extent in L6 (Wang et al., 2013), while projections from high order thalamic nuclei make functional synapses with superficial and deep layers (Sherman, 2012; Audette et al., 2018). In GC, inputs from VPMpc extend to all layers but show laminar-specific differences in the magnitude of evoked responses, with the largest responses in L4. As previously reported for lateral geniculate nucleus (LGN) inputs to L4 of the primary visual cortex (V1), VPMpc stimulation evoked an excitatory response and a large, disynaptic inhibitory response in pyramidal neurons (Kloc and Maffei, 2014), indicating that in both circuits the thalamocortically evoked excitatory/inhibitory ratio is in favor of the latter, thus incoming inputs in both regions are gated by inhibition. However, direct thalamocortical inputs to FS inhibitory neurons in V1 are much larger than those onto pyramidal neurons (Kloc and Maffei, 2014), while in GC they are comparable in amplitude. Furthermore, thalamocortical inputs onto FS neurons in visual, auditory, and somatosensory cortices show more short-term depression than those onto PYR neurons (Reichova and Sherman, 2004; Gabernet et al., 2005; Kloc and Maffei, 2014), as opposed to GC where VPMpc-EPSCs show reduced short-term depression compared to PYR. This evidence suggests that the gating of inputs in GC circuits does not depend necessarily on fast, powerful and short-lasting feedforward inhibition, but rather on a longer lasting feedback inhibitory modulation of incoming activity. This difference in inhibitory modulation between GC and other sensory cortex may carry important functional significance as they may explain the uniquely long-lasting firing dynamics observed for taste responses in GC. The nature of the taste stimulus, which engages complex components including peripheral taste detection by taste receptor cells on the tongue, palate and gums, as well as retronasal activation of receptors in the posterior portion of the mouth and receptors positioned in the early digestive tract (Fujita, 1991; Tizzano et al., 2011; Treesukosol et al., 2011; Samuelsen and Fontanini, 2017; Blankenship et al., 2019; Xi et al., 2022), is likely to engage the taste cortical circuit differently compared to other sensory systems. The distinctive properties of the VPMpc input may endow this thalamocortical system with the ability to support the complex input integration and circuit dynamics characterizing GC’s responsiveness to the gustatory experience.

## Acknowledgments

This work was supported by

National Institutes of Health grant R01DC019827 to AM

National Institutes of Health grant R01DC013770 to AM and AF

National Institutes of Health grant R01DC015234 to AF and AM

National Institutes of Health grant UF1NS115779 to AF and AM

## References

Audette NJ, Urban-Ciecko J, Matsushita M, Barth AL (2018) POm Thalamocortical Input Drives Layer-Specific Microcircuits in Somatosensory Cortex. Cereb Cortex 28:1312–1328.

Blankenship ML, Grigorova M, Katz DB, Maier JX (2019) Retronasal Odor Perception Requires Taste Cortex, but Orthonasal Does Not. Curr Biol 29:62–69 e63.

Cardin JA, Palmer LA, Contreras D (2008) Cellular mechanisms underlying stimulus-dependent gain modulation in primary visual cortex neurons in vivo. Neuron 59:150–160.

Cruikshank SJ, Lewis TJ, Connors BW (2007) Synaptic basis for intense thalamocortical activation of feedforward inhibitory cells in neocortex. Nature Neuroscience 10:462–468.

Cruikshank SJ, Ahmed OJ, Stevens TR, Patrick SL, Gonzalez AN, Elmaleh M, Connors BW (2012) Thalamic control of layer 1 circuits in prefrontal cortex. J Neurosci 32:17813–17823.

Fujita T (1991) Taste cells in the gut and on the tongue. Their common, paraneuronal features. Physiol Behav 49:883–885.

Gabernet L, Jadhav SP, Feldman DE, Carandini M, Scanziani M (2005) Somatosensory Integration Controlled by Dynamic Thalamocortical Feed-Forward Inhibition. Neuron 48:315–327.

Haley MS, Fontanini A, Maffei A (2016) Laminar- and Target-Specific Amygdalar Inputs in Rat Primary Gustatory Cortex. Journal of Neuroscience 36:2623–2637.

Ho J, Tumkaya T, Aryal S, Choi H, Claridge-Chang A (2019) Moving beyond P values: data analysis with estimation graphics. Nature Methods 16:565–566.

Jones LM, Fontanini A, Sadacca BF, Miller P, Katz DB (2007) Natural stimuli evoke dynamic sequences of states in sensory cortical ensembles. Proceedings of the National Academy of Sciences 104:18772–18777.

Katz DB, Simon SA, Nicolelis MAL (2001) Dynamic and Multimodal Responses of Gustatory Cortical Neurons in Awake Rats. Journal of Neuroscience 21:4478–4489.

Kloc M, Maffei A (2014) Target-specific properties of thalamocortical synapses onto layer 4 of mouse primary visual cortex. J Neurosci 34:15455–15465.

Lin JY, Mukherjee N, Bernstein MJ, Katz DB (2021) Perturbation of amygdala-cortical projections reduces ensemble coherence of palatability coding in gustatory cortex. Elife 10.

Liu H, Fontanini A (2015) State Dependency of Chemosensory Coding in the Gustatory Thalamus (VPMpc) of Alert Rats. J Neurosci 35:15479–15491.

Mann EO, Kohl MM, Paulsen O (2009) Distinct Roles of GABAA and GABAB Receptors in Balancing and Terminating Persistent Cortical Activity. Journal of Neuroscience 29:7513–7518.

Olsen SR, Bortone DS, Adesnik H, Scanziani M (2012) Gain control by layer six in cortical circuits of vision. Nature 483:47–52.

Petreanu L, Huber D, Sobczyk A, Svoboda K (2007) Channelrhodopsin-2–assisted circuit mapping of long-range callosal projections. Nature Neuroscience 10:663–668.

Piette CE, Baez-Santiago MA, Reid EE, Katz DB, Moran A (2012) Inactivation of Basolateral Amygdala Specifically Eliminates Palatability-Related Information in Cortical Sensory Responses. Journal of Neuroscience 32:9981–9991.

Pritchard TC, Hamilton RB, Norgren R (1989) Neural coding of gustatory information in the thalamus of Macaca mulatta. J Neurophysiol 61:1–14.

Reichova I, Sherman SM (2004) Somatosensory corticothalamic projections: distinguishing drivers from modulators. J Neurophysiol 92:2185–2197.

Ruiz A, Campanac E, Scott RS, Rusakov DA, Kullmann DM (2010) Presynaptic GABAA receptors enhance transmission and LTP induction at hippocampal mossy fiber synapses. Nat Neurosci 13:431–438.

Samuelsen CL, Fontanini A (2017) Processing of Intraoral Olfactory and Gustatory Signals in the Gustatory Cortex of Awake Rats. J Neurosci 37:244–257.

Samuelsen CL, Gardner MPH, Fontanini A (2012) Effects of Cue-Triggered Expectation on Cortical Processing of Taste. Neuron 74:410–422.

Samuelsen CL, Gardner MPH, Fontanini A (2013) Thalamic Contribution to Cortical Processing of Taste and Expectation. Journal of Neuroscience 33:1815–1827.

Sanchez-Vives MV, Barbero-Castillo A, Perez-Zabalza M, Reig R (2021) GABAB receptors: modulation of thalamocortical dynamics and synaptic plasticity. Neuroscience 456:131–142.

Sherman SM (2012) Thalamocortical interactions. Curr Opin Neurobiol 22:575–579.

Sherman SM, Guillery RW (1998) On the actions that one nerve cell can have on another: distinguishing “drivers” from “modulators”. Proc Natl Acad Sci U S A 95:7121–7126.

Stone ME, Fontanini A, Maffei A (2020) Synaptic Integration of Thalamic and Limbic Inputs in Rodent Gustatory Cortex. eNeuro 7:ENEURO.0199-0119.2019.

Tan Z, Hu H, Huang ZJ, Agmon A (2008) Robust but delayed thalamocortical activation of dendritic-targeting inhibitory interneurons. Proc Natl Acad Sci U S A 105:2187–2192.

Tizzano M, Cristofoletti M, Sbarbati A, Finger TE (2011) Expression of taste receptors in solitary chemosensory cells of rodent airways. BMC Pulm Med 11:3.

Treesukosol Y, Smith KR, Spector AC (2011) The functional role of the T1R family of receptors in sweet taste and feeding. Physiol Behav 105:14–26.

Verhagen JV, Giza BK, Scott TR (2003) Responses to taste stimulation in the ventroposteromedial nucleus of the thalamus in rats. J Neurophysiol 89:265–275.

Wang L, Kloc M, Gu Y, Ge S, Maffei A (2013) Layer-specific experience-dependent rewiring of thalamocortical circuits. J Neurosci 33:4181–4191.

Wang L, Kloc M, Maher E, Erisir A, Maffei A (2019) Presynaptic GABAA Receptors Modulate Thalamocortical Inputs in Layer 4 of Rat V1. Cereb Cortex 29:921–936.

Wehr M, Zador AM (2003) Balanced inhibition underlies tuning and sharpens spike timing in auditory cortex. Nature 426:442–446.

Xi R, Zheng X, Tizzano M (2022) Role of Taste Receptors in Innate Immunity and Oral Health. J Dent Res 101:759–768.

Yamamoto K, Koyanagi Y, Koshikawa N, Kobayashi M (2010) Postsynaptic Cell Type–Dependent Cholinergic Regulation of GABAergic Synaptic Transmission in Rat Insular Cortex. Journal of Neurophysiology 104:1933–1945.

